# Fast reoptimization of human motor patterns in non-Earth gravity fields locally induced by a robotic exoskeleton

**DOI:** 10.1101/2022.11.10.516038

**Authors:** Dorian Verdel, Simon Bastide, Franck Geffard, Olivier Bruneau, Nicolas Vignais, Bastien Berret

## Abstract

Gravity is a ubiquitous component of our environment that we learnt to optimally integrate in movement control. Yet, altered gravity conditions arise in numerous applications from space exploration to rehabilitation, thereby pressing the sensorimotor system to adapt. Here, we used a robotic exoskeleton to test whether humans can quickly reoptimize their motor patterns in arbitrary gravity fields, ranging from 1g to −1g and passing through Mars- and Moon-like gravities. By comparing the motor patterns of actual arm movements with those predicted by an optimal control model, we show that our participants (N = 61) quickly and optimally adapted their motor patterns to each local gravity condition. These findings show that arbitrary gravity-like fields can be efficiently apprehended by humans, thus opening new perspectives in arm weight support training in manipulation tasks, whether it be for patients or astronauts.

## Introduction

Earth’s gravity has pervasive effects on human neuromechanics and motor control. Several studies have suggested that our central nervous system (CNS) has an internal representation of gravity, spread over different brain areas, which allows to optimize the control of movement with respect to the ambient gravity field (*1–4*). The most direct evidence of such a gravity-exploitation theory came from studies conducted with astronauts in–or returning from– missions and during parabolic flights (*5–13*). Indirect evidence was also obtained by comparing the characteristics of vertical and horizontal movements (*9, 10, 14–20*). In particular, several studies reported consistent kinematic differences between vertical and horizontal movements, which gradually vanished through the adaptation to microgravity or could be recreated when applying a gravity-like force field in microgravity (*6,21*). This evidence was further supported by congruent observations of muscle patterns, which were found to substantially differ depending on motion direction with respect to gravity, in both humans and monkeys (*20, 22–24*). Importantly, the adaptation of kinematic and muscular patterns to the ambient gravity field was found to comply with the predictions of optimal control models based on effort minimization. Several model based studies supported the hypothesis that the CNS optimally exploits gravity during arm motor planning (*5,9,20,25–28*).

Interestingly, the gravity-exploitation theory makes specific predictions in arbitrary gravity fields which have been untested so far. Studies in parabolic flights have allowed to test this theory in a couple of hypo- and hyper-gravity fields with a limited number of trials and participants (*5, 9, 12, 13*). Moreover, while very relevant to space exploration (*29–33*), completely immersing participants in a novel gravity field is not representative of other applications. For instance, it is common to use devices to support a patient’s limb and reduce their muscle effort required to counteract gravity in rehabilitation (*34–42*). In this case, the neuromechanical system is locally impacted, mostly through somatosensory information. In principle, this information could be sufficient to trigger an optimal motor adaptation to the new mechanical context that alters the human’s gravitational torque at the joints. Indeed, the somatosensory system has been shown to play a predominant role for learning new dynamics efficiently (*43,44*). In particular, in macaque monkeys, proprioception was shown to provide sufficient information to accurately perceive gravity, within three weeks after bilateral vestibular loss (*45*). Alternatively, it is also possible that congruent information from other sensory systems (e.g. visual, proprioceptive, vestibular) is required to update the internal representation of gravity and trigger an optimal adaptation (*7,13,14,46,47*). A merging of different sensory cues might thus be necessary to optimally exploit gravity in such a novel environment.

To test whether participants can reoptimize their motor patterns in arbitrary local gravity fields, the present paper leverages recent advances regarding arm weight compensation with active exoskeletons (*48*) to induce gravity conditions that were hitherto hardly achievable. This follows a general path towards the exploitation of robotics in neuroscience (*49*). Here we used the ABLE robotic exoskeleton (*50*) to induce various local gravity fields, ranging from normal gravity (1g) to reversed gravity (−1g) and passing through microgravity (0g). A gradual change of gravity with a 0.2g-step ranging from −1g to 1g was also applied on the forearm of human participants, thereby including Mars- and Moon-like gravities. We then compared experimental results to the predictions of the gravity-exploitation theory. The predictions of a representative model, termed Smooth-Effort (SE) model (*20*), are illustrated for upward movements in Figure 1. The changes in velocity profiles are quantified in terms of the relative time to peak velocity (rtPV), which is known to be a robust gravity-dependent parameter. This gravity-dependent asymmetry of velocity profiles for single-joint movements is well documented and is predicted to vary in the 1g, 0g and −1g conditions if local gravity is exploited. Furthermore, a striking non-linear evolution of rtPV is even predicted when varying gravity more finely because the exploitation of gravity depends on its actual effect on the limb’s acceleration for a fixed movement duration. If these theoretical predictions match experimental data, this would reinforce the gravity-exploitation theory and indicate that somatosensory information is sufficient to allow humans reoptimizing their motor patterns according to the locally induced gravity. If theoretical predictions do not match experimental data, then either incongruent sensory signals prevent the reoptimization of movement with respect to the local gravity or the theoretical model must be revised.

**Figure 1:**
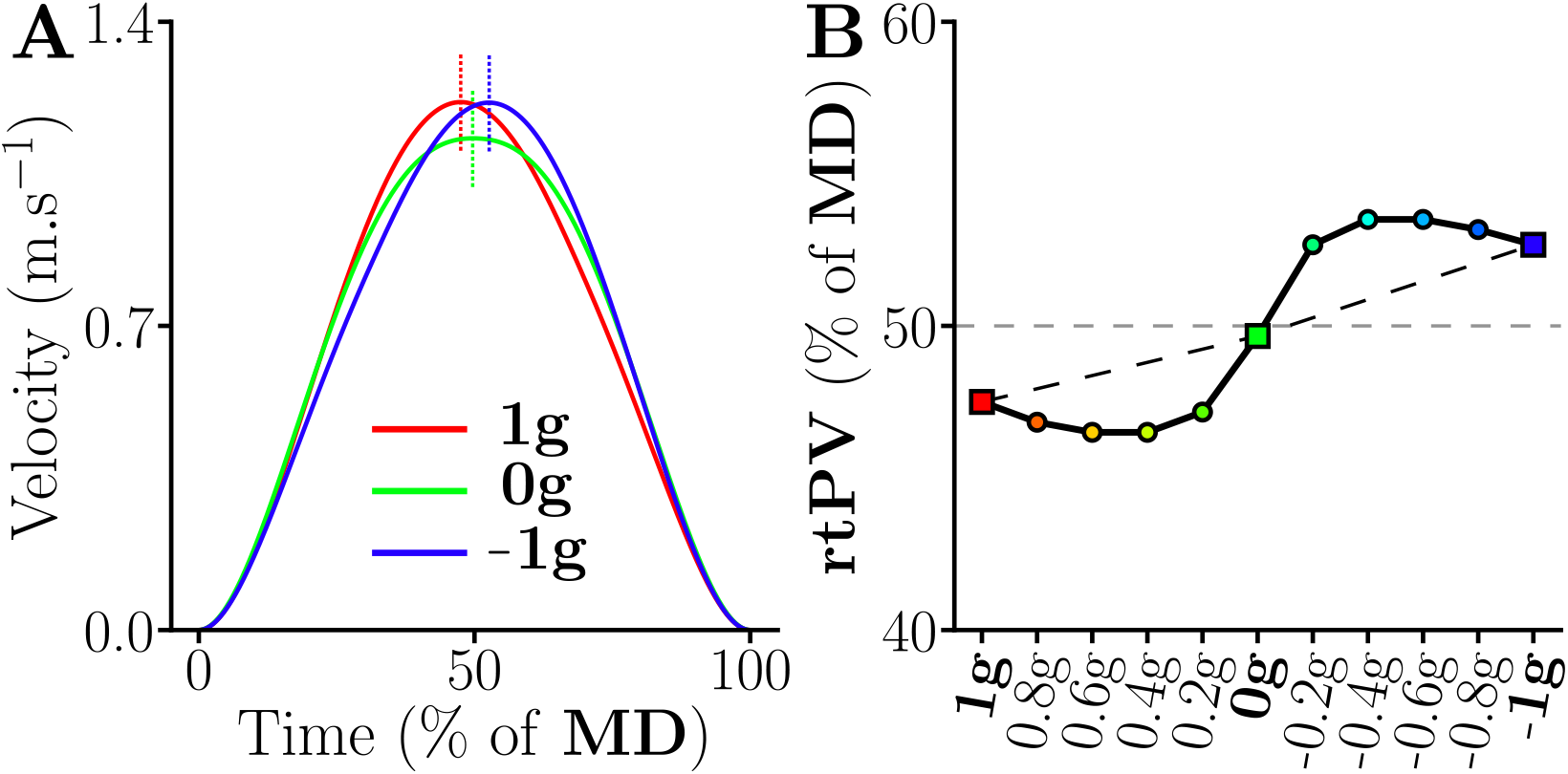
Predictions of the gravity-exploitation theory according to the Smooth-Effort (SE) optimal control model of (*20*). We simulated fast upward pointing movements with the forearm of duration MD=0.6 s, while only varying gravity. **A**. Predicted velocity profiles for the 1g, 0g and −1g conditions. Vertical dashed lines indicate where the peak velocity (PV) is reached, which highlights the variations of the relative time-to-peak-velocity parameter (rtPV) according to the model. **B**. Predicted evolution of rtPV (in percentage of movement duration, MD) for 11 gravity fields ranging between 1g and −1g. While a linear gradient can be seen between the 1g, 0g and −1g conditions, the finer-grained analysis reveals a non-linear evolution of rtPV across gravities.

## Results

We asked *N* = 61 participants to perform fast upward pointing movements of 45° with the forearm toward semi-spherical targets of radius 2 cm. The participants were connected to a robotic exoskeleton that was controlled to mechanically emulate various gravitational torques at the participant’s elbow joint. The control law was individualized thanks to a thorough identification procedure conducted prior to each experiment and validated in a previous study (*48*). The task is illustrated in Fig. 2A and the reader is deferred to the Material and Methods for more details about the experimental procedures.

**Figure 2:**
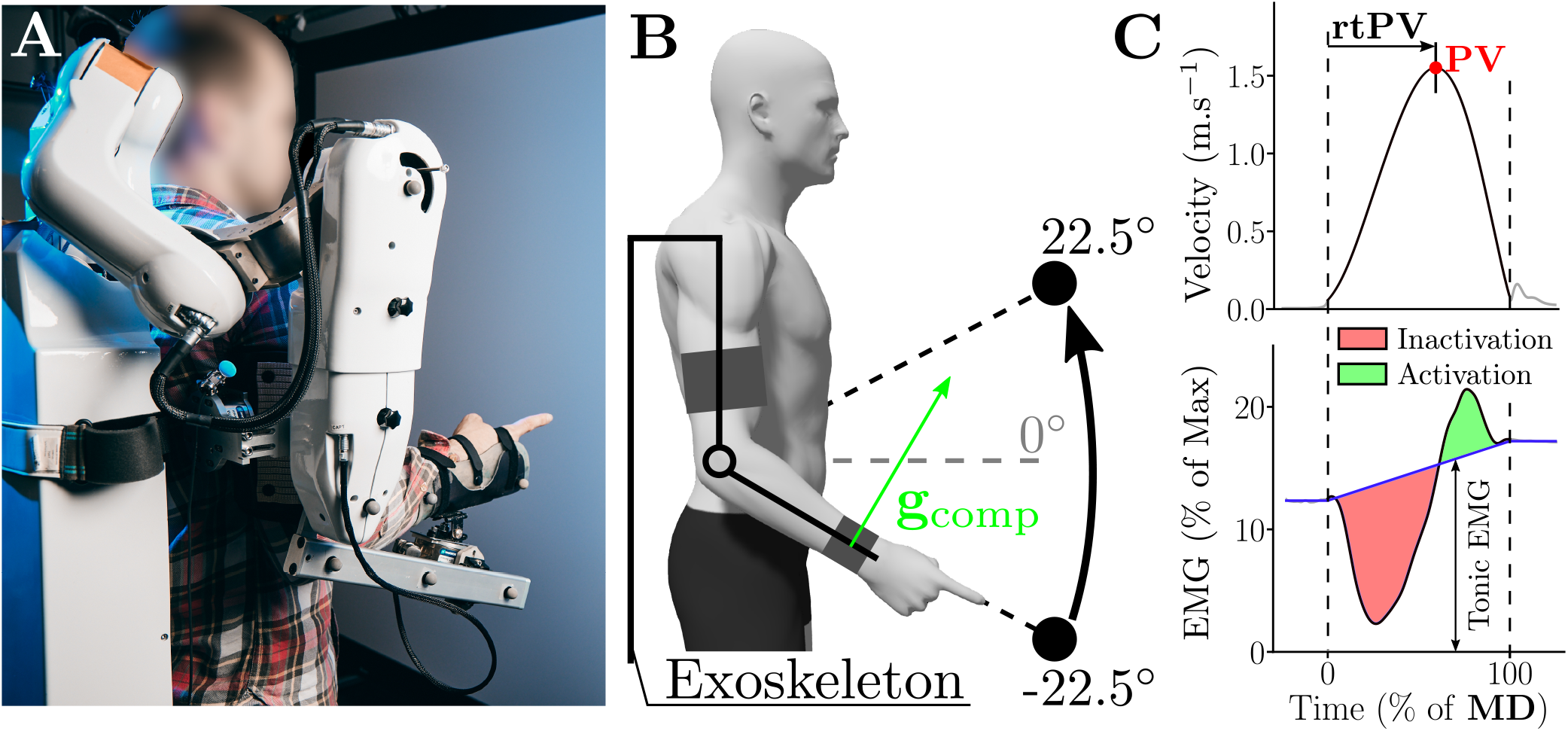
**A**. Illustration of a participant connected to the exoskeleton during the pointing task. **B**. Schematic illustration of the motor task, consisting in upward forearm movements (here of 45°) with the human connected to the ABLE robotic exoskeleton. A force/torque sensor placed at the interface between the participant and the exoskeleton (about the wrist joint) allowed tracking a desired normal force g*_comp_*, mechanically corresponding to the effect of gravity fields ranging from 1g to −1g. The real-time forearm inclination and joint misalignments between the human and the robot were taken into account to accurately estimate g*_comp_* (*48*). **C**. Definition of the main kinematic and EMG parameters analyzed in the present study for an example movement in the −1g condition. Fast pointing movements are typically characterized by velocity profiles that mainly consist of one acceleration phase and one deceleration phase. Here, the temporal structure of movement was characterized by the rtPV parameter, as it is known to be sensitive to gravity (*9*). The represented EMG data are the envelope of the filtered, rectified and normalized signal recorded during −1g motions. Regarding EMG data, we systematically subtracted the tonic activity (blue line) from rectified EMGs and analyzed the resulting phasic EMG patterns. This phasic activity is also known to be sensitive to gravity (*20*). Negative phasic activity is referred to as inactivation and indicates periods when the EMG activity of a muscle is below the tonic level that would be required to counteract gravity (in shaded red). Positive phasic activity, referred to as activation (in shaded green), is responsible for net accelerations in the gravity field.

A series of three experiments was conducted in this work. In the first two experiments, three gravity fields (1g, 0g, −1g) were tested to analyze if and how participants changed their motor patterns according to the locally induced gravity fields (Experiment 1 and Experiment 2). In the last experiment, a gradient of 11 gravity fields, ranging from −1g to 1g with a step of 0.2g was tested to assess whether gradual changes of local gravity lead to nonlinear changes of motor patterns in agreement with the gravity-exploitation theory. The main kinematic and muscular parameters used to quantify motor patterns are defined in Fig. 2B.

### Motor patterns change quickly and significantly with respect to the local gravity field

In Experiment 1, participants (*N* = 22) performed 6 consecutive blocks of fifteen 45-degrees upward movements in each of the 1g, 0g, −1g conditions. The order of the conditions was randomized. Analyses conducted on the first 10 movements of participants involved in Experiment 1 allowed to exhibit that a quick adaptation occurred during the very first movements after which a plateau was reached and the participants could reliably execute the task. In 0g and −1g the main kinematic parameters did not differ from the grand-average value computed from the last five blocks, which means that a plateau was attained after a couple of movements (see supplementary Fig. S.1). The same observation was made for the main EMG parameters (see supplementary Fig. S.2). No evidence of a longer term adaptation was found across the 6 blocks performed in each condition of Experiment 1, for all the tested kinematic and EMG parameters (see supplementary Fig. S.3). Furthermore, no adaptation across blocks was found on parameters usually impacted by gravity (see supplementary Fig. S.4). As a consequence, all the subsequent analyses were conducted on data averaged across all the blocks (excluding the first 5 trials of block 1), and we focus hereafter on the changes induced by different gravity conditions on the average motor patterns.

#### Kinematic analysis

Figure 3 depicts the average motor patterns (position and velocity, and phasic EMGs) for a representative participant in the 3 gravity conditions of Experiment 1.

**Figure 3:**
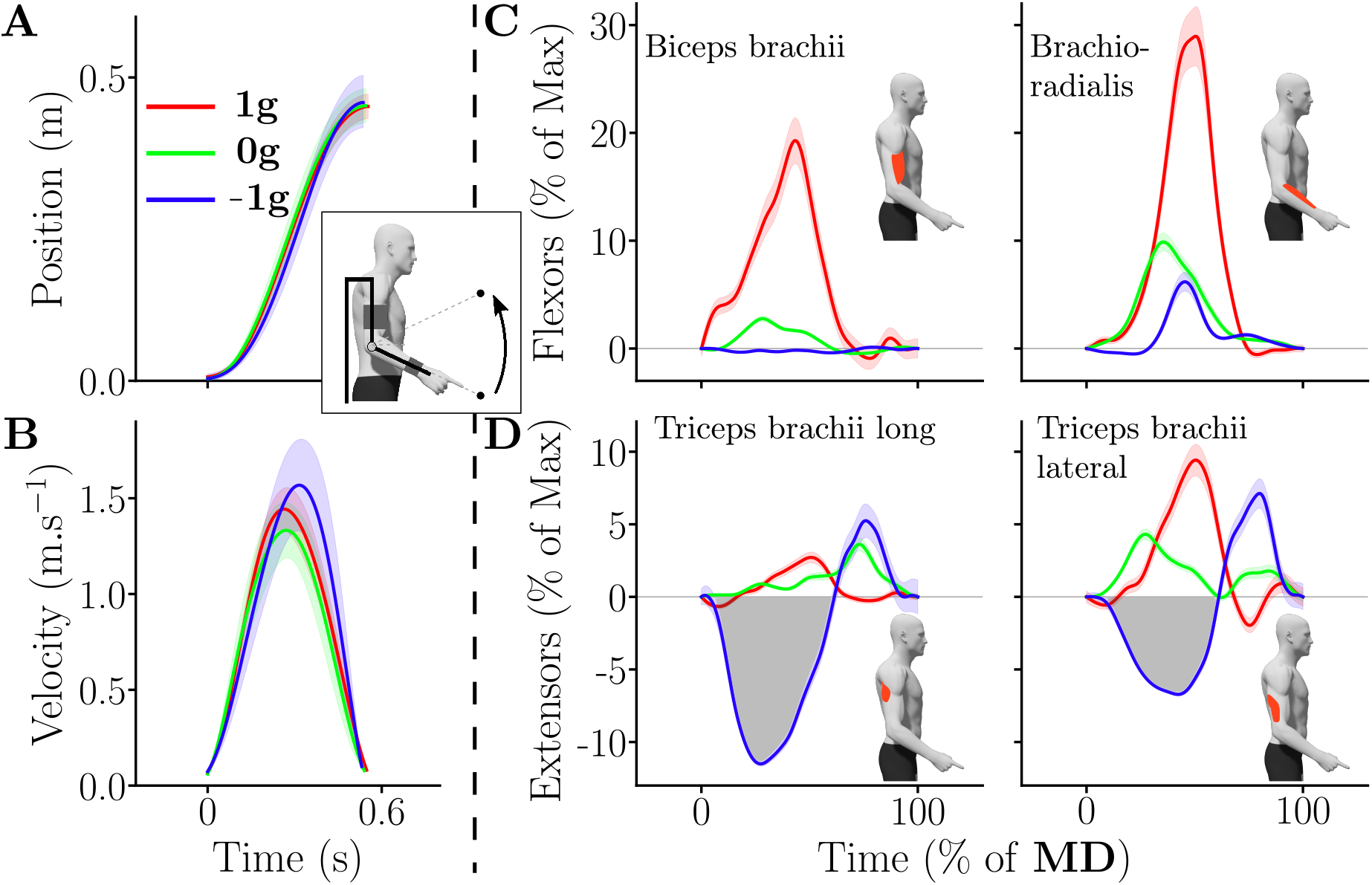
Kinematics and phasic EMG envelopes for one representative participant in the different gravity conditions (red=1g, green=0g and blue=−1g). **A**. Mean hand displacement. Standard errors across trials are represented as shaded areas. **B**. Mean hand velocity profile. **C,D**. Mean phasic flexors and extensors muscle patterns in the different gravity conditions, normalized in time (MD was about 0.6 s). Notably, an early inactivation of the extensors (negative phasic area, emphasized by the grey shaded area) in the −1g condition can be observed (see blue trace in panel **D**).

As already mentioned, an interesting parameter theoretically revealing gravity exploitation and sensitive to the ambient gravity acceleration is rtPV (*9*). This parameter allows comparing kinematic motor patterns of movements of different durations. Here, only slight variations of durations and speeds were found between gravity conditions. Movements performed in −1g tended to be slightly faster than those performed in 0g and 1g, as reflected by a repeated-measures ANOVA conducted on MD (*p* = 0.004, 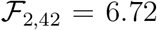 and *η*^2^ = 0.24), but no difference on MD was found across conditions with post-hoc pairwise comparisons (*p* > 0.18 for all comparisons). Regarding rtPV, group and individual data are depicted in Fig. 4A, revealing a main effect of the local gravity field on rtPV (repeated-measures ANOVA: *p* < 0.001, 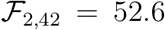 and *η*^2^ = 0.71). Post-hoc analyses indicated that the rtPV was significantly higher in the 0g and −1g conditions when compared with the 1g condition (*p* < 0.002 in both cases). This means that velocity profiles were more left-skewed in 0g and −1g than in 1g. However, no difference was found between the 0g and the −1g conditions in this sample (*p* = 0.225).

**Figure 4:**
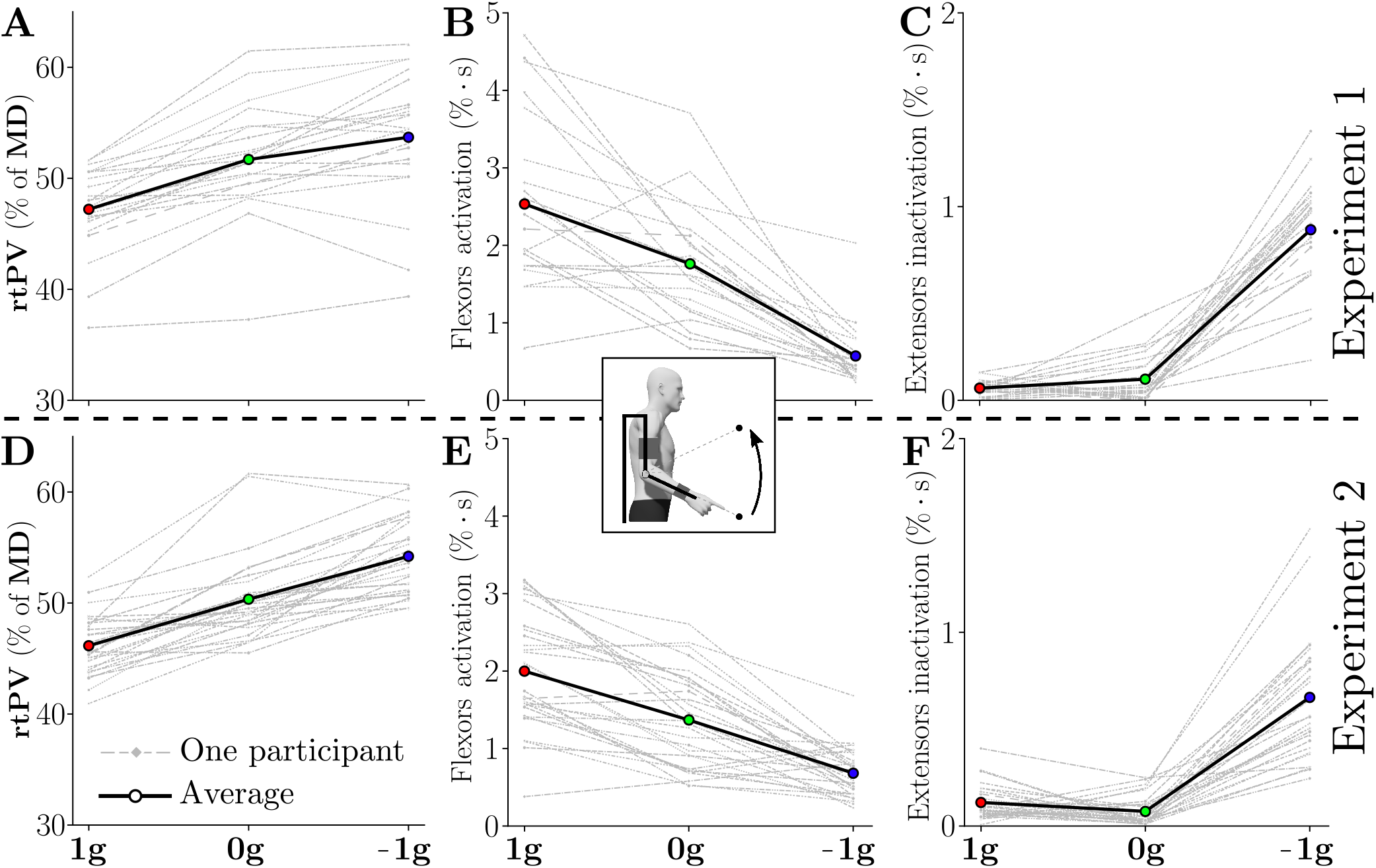
Average kinematic and muscular behavior of each participant (grey) and of the tested population (black bold) for both Experiment 1 and Experiment 2. **A,D**. Adaptation of rtPV across conditions. **B,E**. Adaptation of the flexors activation across conditions. **C,F**. Adaptation of the extensors inactivation across conditions.

To self-replicate our findings and test whether significance can be reached for comparisons between 0g and −1g, we performed an additional experiment including two blocks of 25 upward movements of 45° for each gravity condition, knowing that adaptation was very quick (referred to as Experiment 2, *N* = 29 participants). The rtPV values obtained in this experiment are depicted in Fig. 4D.

The same trends as in Experiment 1 were observed. Repeated-measures ANOVA confirmed a main effect of local gravity on rtPV (*p* < 0.001, 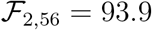 and *η*^2^ = 0.77). Post-hoc tests further revealed significant differences between all gravity conditions in this sample (*p* < 0.001 in all cases).

Together, the results of Experiment 1 and Experiment 2 clearly show that the averaged velocity profiles were robustly tuned according to the local gravity field created by the exoskeleton.

To better understand the cause of this gravity-dependent tuning of velocity profiles, we then investigated the underlying muscular strategies. The patterns of activation of flexor and extensor muscles may reveal the extent to which participants took advantage of the novel gravity field (*20,22–24*).

#### Muscular analysis

Forearm movements are controlled by opposing muscle groups that can be gathered as follows: 1) flexors (e.g. mainly the biceps-brachii and brachio-radialis) and 2) extensors (e.g. mainly the triceps-brachii long and lateral heads). In the Earth’s gravity field, the flexors are antigravity muscles in the sense that they generally allow counteracting the action of gravity. The extensors play the opposite role in 1g and could be termed gravity muscles. In 0g this distinction is irrelevant whereas in −1g the role played by flexors and extensors should be inverted such that extensors should become antigravity muscles.

Qualitatively, the EMG patterns are depicted in Figure 3C,D for each gravity condition for a representative participant. In the 1g condition, the movement was triggered by a strong phasic activation of the flexors. In 0g, the EMG patterns were similarly structured as in 1g, except that the activation of the flexors was much weaker. Finally, the EMG pattern in the −1g condition was very different and started with an inactivation of the extensors, meaning that the participant let the local gravity field created by the robot initiate the upward movement. This negative phasic EMG was followed by an activation of the extensors for decelerating the movement at the end. This negativity existed from the first trial even though its duration was adapted during the first trials (see Fig. S.2).

These qualitative observations were supported by quantitative analyses performed on the activation of flexors and the inactivation of extensors in the different conditions (see Fig. 4B,C,E,F). As for the kinematic analysis, both group and individual data are reported. There were clear differences between the different gravity conditions in terms of flexors activation (see Fig. 4B,E; main effect, *p* < 0.0041, 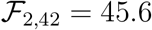 and *η*^2^ = 0.68). Post-hoc analyses indicated that all gravity conditions were different (*p* < 0.001). In particular, there was a 30% decrease in flexors activation between the 1g and 0g conditions and a 68% decrease in flexors activation between the 0g and −1g conditions during Experiment 1. The same significant decrease of flexors activation was observed during Experiment 2 between the three tested conditions (main effect: *p* < 0.0041, 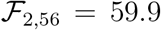 and η^2^ = 0.68; post-hoc: p < 0.003 in all cases).

The inactivation of extensors also exhibited clear differences between the conditions (see Fig. 4C,F; main effect: p < 0.001, 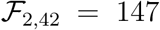 and *η*^2^ = 0.87). In particular, during Experiment 1, the −1g condition was the only one that exhibited clear extensors inactivation, which was reflected in post-hoc comparisons with the 1g and 0g conditions (*p* < 0.001 in both cases). The same trend was observed during Experiment 2 (main effect: *p* < 0.001, 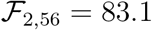 and *η*^2^ = 0.75). The −1g condition was again the only one exhibiting extensors inactivation (post-hoc: *p* < 0.001 in both cases).

In summary, these first two experiments revealed that the participants quickly reoptimized their motor patterns toward effort minimization according to the local gravity fields, in agreement with the prediction of the gravity-exploitation theory. Indeed, the skewness of velocity profiles changed significantly in the 0g and −1g conditions compared to the 1g condition as predicted by the SE model, and coherent changes in muscle activation and inactivation patterns were observed.

To further test the striking prediction of a non-linear evolution of rtPV when gravity is varying more finely, we conducted a third experiment where we gradually varied the gravity compensation with a 0.2g-step.

### Adaptation to gradual changes of gravity and comparisons with optimal control models

In Experiment 3 (*N* = 10 participants), we considered the same 45-degrees upward forearm movements for a gradient of 11 gravity fields, equally spaced between 1g and −1g, and passing through 0g, Mars-like gravity (about 3.7 m.s^-2^, which is around 0.4g) and Moon-like gravity (about 1.6 m.s^-2^, which is slightly under 0.2g).

Results for rtPV are depicted in Fig. 5A. Data showed that rtPV globally tended to increase from 1g to −1g, hence confirming our previous observations. This gradual increase exhibited a non-linear trend on average. The average rtPV varied according to a “sigmoidal” form across the different gravity conditions, with a minimum observed in 0.8g and a maximum observed in −0.6g. Furthermore, between 1g and 0.2g the average rtPV followed a smooth gradient but stayed clearly under 50% of MD, which implies that the acceleration phase was shorter than the deceleration phase (right-skewed velocity profiles). The average rtPV observed in 0g and −0.2g was close to 50% of MD, the acceleration and deceleration phases were therefore almost equivalent (approximately symmetric velocity profiles). Finally, between −0.4g and −1g, the rtPV was above 50% of MD, which implies that the acceleration phase was on average longer than the deceleration phase (left-skewed velocity profiles).

**Figure 5:**
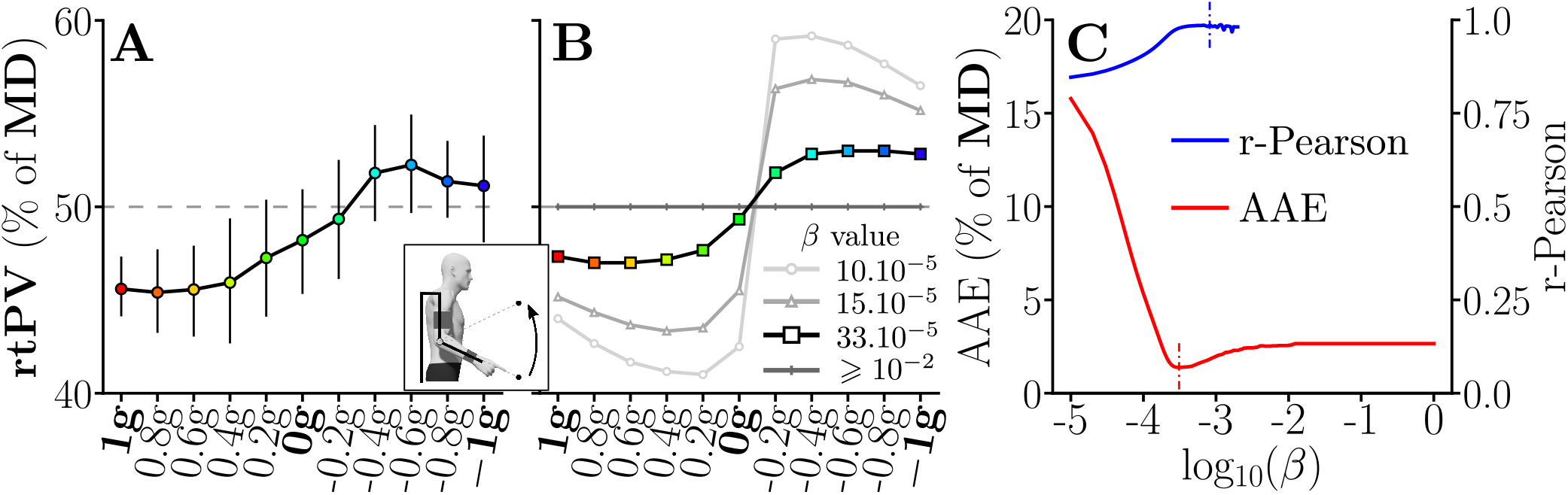
Averaged rtPV adaptation and simulations. **A**. Experimental results of Experiment 3, averaged data are represented with bars representing the standard error. **B**. Simulation results for various weighting of the cost function. The highlighted simulation result corresponds to the optimal weighting of the cost function in terms of average absolute error (AAE). **C**. Evolution of the AAE and of the Pearson correlation coefficient with regard to the cost function weighting with *β* ∈ [10^_5^,1]. The Pearson correlation coefficient was only computed when the predicted rtPV presented enough variability across conditions (not defined otherwise) and plotted for significant correlations (*p* < 0.05). The optimal values of *β* with regard to both these error criteria are highlighted with small dotted bars. **B,C**. These graphs exhibit the evolution of the prediction quality both in terms of gradient and average value with regard to the simulated compromise between smoothness and effort.

This evolution is reminiscent of the one predicted by the SE model given in Introduction (see Fig. 1B), representative of the gravity-exploitation theory (*9,20*). This model allows simulating movements under the assumption of the minimization of a compromise between the absolute work of muscle torques and the time-integral of squared jerk, which corresponds to an effort minimization model with smoothness regularization. In its present form, the model had one free parameter in the cost function, which sets the compromise between the effort and smoothness terms (*β* in Eq.4). Predictions of the SE model in terms of rtPV have already been exemplified in Fig. 1A, which can be compared qualitatively to 5B. Here, the quality of the model prediction was assessed more quantitatively by computing the average absolute error (AAE) and the Pearson correlation coefficient between the experimental and simulated rtPVs. The goal was to evaluate the extent to which the gravity-dependent tuning of this parameter was captured by the model.

Furthermore, by varying *β*, we could thus search for an optimal weighting of the cost function to match the experimental data, which corresponds to an inverse optimal control approach (e.g. identifying the weighted cost function that best fits experimental data). The results of this procedure are depicted in Figures 5B,C. The SE model was able to provide both small estimation errors on rtPV (AAE ≈ 1.38% of MD with an optimal *β* = 3.3 × 10^-4^) and high Pearson correlation coefficients (*r*-Pearson > 0.985 with an optimal *β* = 8.2 × 10^-4^). Models with too small or too large *β* either predicted overestimated or underestimated changes of rtPV with respect to the local gravity field. Interestingly, these values of *β* were close to those reported in previous works to predict the kinematics of one degree of freedom shoulder movements (*9*), which shows a certain consistency of the model predictions across different experimental paradigms.

## Discussion

In the present paper, we tested the gravity-exploitation theory by investigating if and how the humans can reoptimize their motor patterns in arbitrary gravity fields locally induced by a robotic exoskeleton. Both kinematic and muscle patterns were analyzed during upward pointing movements involving the forearm while the exoskeleton applied various gravity fields like 0g (weightlessness), 0.4g (close to Mars’ gravity) or −1g (inverted Earth’s gravity). In a series of three experiments, we found that participants quickly changed their motor patterns according to the local gravity, and tended to take advantage of it whenever possible. Indeed, varying gravity from 1g to −1g was concomitant with a shift of hand velocity profiles from right-skewed to left-skewed. Furthermore, EMG patterns were in line with an exploitation of the gravity direction and magnitude to accelerate the limb. This was exemplified by the fact that upward movements systematically started with the inactivation of the extensors in −1g. These empirical observations were in good agreement with the predictions of an optimal control model based on effort minimization, thereby providing computational evidence for the gravity-exploitation theory. Below, we discuss these findings with respect to the internal model of gravity that could allow participants to quickly reoptimize their motor patterns in such novel gravity fields from somatosensory information.

Our participants quickly changed their motor patterns in 0g and −1g. Plateaus on all the main kinematic parameters were reached after a couple of trials. Interestingly, plateaus were reached instantaneously for most parameters computed on muscle activities. A similar conclusion had been drawn with the same exoskeleton when increasing the apparent inertial torque without affecting the participants’ gravitational torque (*51*). However, such a quick adaptation contrasts with other works showing a slower adaptation to microgravity for similar single-joint arm movements, which took about 75 movements during parabolic flights to completely remove up/down differences in velocity profiles (*9*). The main difference between these studies seems to lie in the nature of the sensory changes induced by the novel gravito-inertial environment. During parabolic or space flights, the sensory changes are global and captured by both the vestibular and somatosensory systems (*7, 13*), which has been suggested to require learning before a possible adaptation (*52, 53*). In the present human-exoskeleton interaction study, only the somatosensory signals of the assisted limb were affected, hence inducing only a local sensory change. Our findings suggest that motor planning may be more efficiently adapted in this case, despite incongruent gravity-related sensory cues. These results are consistent with previous studies performed with different experimental paradigms, which suggested that somatosensory feedback, in particular haptic, allows a fast and significant adaptation (*11, 54–56*). This is probably due to the very quick adaptation of somatosensory loops and to the predominant role of the somatosensory system in the early learning of new dynamics (*43, 44, 57, 58*). Such a quick adaptation could therefore be a feature of the CNS to easily handle object manipulation, even when deprived of specific visual or vestibular information (*59*), which is a common type of action in daily life. Here, the gravity torques induced by robot essentially induced a parametric change in the human limb’s dynamics (*60*), which could be quickly estimated from somatosensory cues even in statics as in previous works (*61*). Broadly affecting all sensory systems at once via a global gravity change may thus involve more complex processes in the CNS. For example, a potential reweighing of multiple sensory inputs may take longer to trigger a fine reoptimization of movement (*9,62,63*).

Here, the adaptation was not only quick but also exhibited the signature of effort-based optimality in the local gravity field. The asymmetry of velocity profiles (e.g. rtPV) was shown to change consistently across the different gravity conditions in all three experiments. The relative duration of the acceleration phase tended to increase when the induced gravity decreased from 1g to −1g and this evolution was not simply linear. All these changes were well replicated by the Smooth-Effort optimal control model (see Fig. 5A–C). For example, the maximum rtPV was reached around the −0.6g condition for most participants and then it started to decrease (see Fig. 5A), in agreement with model predictions. Furthermore, when gravity was reversed and pushed the participant’s forearm upward, longer relative duration of acceleration was observed, thereby increasing the rtPV parameter. This particular kinematic strategy has been observed previously during downward motions in the Earth’s gravity field (*5,9,10,16–20*) and was well explained by the gravity-exploitation theory (*9,20,25*). In particular, from the condition −0.4g, the average rtPV exceeds 50% of MD in Experiment 3. Interestingly, this strategy contrasts with accuracy constraints typically associated with rapid movements as it is known that deceleration is instead larger when participants are required to be maximally fast and accurate (*64*). Therefore, the left-skewness of velocity profiles was clearly not due to accuracy concerns but proved to be compatible with an optimal movement strategy taking into account the current gravity acceleration to minimize effort.

At the muscle level, changes in activation and inactivation patterns were also compatible with an exploitation of the local gravity (e.g. 1g, 0g and −1g conditions). The most notable effect is likely the very quick adaptation of extensors to the −1g condition. In this condition, the movement was initiated with the inactivation of the extensors rather than with the activation of the flexors (see Fig. 4C,D). This behavior is particularly revealing of the human capacity to optimally integrate the local gravity field. Even though an inverted gravity field is very unusual, the participants were all able to take advantage of its presence to accelerate their limb with low effort (see Fig. 4D,H). In the Earth’s gravity field, this inactivation pattern is usually observed on the flexors, to use gravity to accelerate downward movements at their initiation (*20, 23, 24*). All the participants were able to quickly assign to their extensors a role normally assigned to their flexors. This was probably possible thanks to the somatosensory information gathered in statics before movement execution.

Overall, our experimental and computational analyses agree with the gravity-exploitation theory and not with an alternative gravity-compensation theory, which would predict that gravity should not affect kinematic motor patterns. In this case, the phasic activity would only reflect inertia-related efforts while tonic activity would always be present to counteract gravity-related efforts (*16,65,66*). In our study, it was clear that the participants did not attempt to counteract the force applied by the exoskeleton and maintain an unchanged kinematic strategy for all local gravity fields. The observed changes of motor patterns were not arbitrary either, as they could have been if the system had failed to account for the new gravities. Instead, our results point to a specific and optimal-like adaptation to each local gravity field.

It is worth stressing that different models could have been used as representatives of the gravity exploitation theory. Here, we used a previously proposed model based on the minimization of the absolute work of muscle torques (*25*). We did not consider complex muscle dynamics to reduce the number of unknown parameters and simplify the modeling. Hence, only one parameter was left free in the model, which served to set the compromise between effort and smoothness optimization. Using an inverse optimal control approach (*26*), we identified the best-fitting *β* coefficient to replicate the data, which was relatively close to that proposed in previous studies. This analysis revealed that minimizing effort too strongly tended to overestimate the effects of gravity on the kinematics whereas maximizing smoothness tended to underestimate them. It could be interesting in the future to investigate the effects of using more accurate muscle models to refine the estimation of energy expenditure in such tasks (*67*). Nevertheless, conducting a sensitivity analysis related to the uncertainty about the various model parameters could be a tedious task for such optimal control simulations. Here, because we mainly wanted to examine how gravity can be exploited in motor planning, we examined a simpler model that cannot impute the gravity-dependent kinematic changes to the low-level contraction properties of muscles, in accordance with previous interpretations (*9, 14, 17, 20*). Yet, studying longer term adaptations to altered gravity environment could provide an interesting avenue for research because major neuromuscular reorganizations may then occur (*68*), which could strongly affect motor patterns. Alternatively, estimating metabolic energy through gas exchanges like in other studies (*69, 70*) could be interesting to estimate the different costs of movement in the various gravity environments and check whether a minimum is attained in the 0g condition for instance, or whether the costs decrease during longer-term adaptations.

Our current conclusions were obtained with a robotic exoskeleton controlled in such a way that arbitrary gravity force field could be generated at the level of the human joint. Although a subject specific calibration was conducted to create the desired local gravity as accurately as possible, there are currently two limits that should be mentioned. The first one is that exoskeletons may disturb the human motor behavior even in transparent mode (our 1g condition here), although interaction efforts are minimized. We have quantified the nature of the perturbation for elbow flexions/extensions in previous works and the remaining disturbances were mainly due to the additional inertia caused by the exoskeleton (*48, 71–73*). It could be possible to compensate this inertia by using predictive control methods (*74*). However, the inherent variability of human movement makes difficult the design of a simple and straightforward compensation method for human-exoskeleton interaction. One alternative, on which the present paper is based, is to assess the nature of the perturbation introduced by the exoskeleton *a posteriori* and integrate it in the models as an augmented inertia for the coupled human robot system. The second limitation lies in the tracking of the desired gravity field. Residual errors will inevitably occur due to inherent limits of the control law. These errors were quantified in a previous paper (*48*). Although relatively small, slight variations from a true gravitational filed are present and it is difficult to estimate the extent to which it could affect the participants’ behavior. However, substantial changes in the rtPV parameter were observed between gravity fields. It is possible that the behavior of the robot could slightly alter what would be observed after steady adaptation to the real Moon’s or Mars’ environments. However, the ease with which gravity can be locally changed, even approximately, compared to space missions, may nevertheless be an appealing way to test how the CNS adapt to novel gravito-inertial environments.

Finally, beyond gaining fundamental knowledge, understanding how the human CNS integrates gravity has numerous applications. In particular, this question is critical to space exploration with astronauts (*63, 75–78*). There, the use of active exoskeletons could be envisioned during long-term missions as a way to limit muscle atrophy (*79–81*) (recreating Earth’s gravity in spaceships as a countermeasure) or as a way to train astronauts to perform dexterous manipulation tasks in various gravity environments (*6, 29*). Furthermore, weight compensation for patients suffering from muscle weakness is an appealing approach in neurorehabilitation. For example, post-stroke patients can recover more mobility when gravity-related efforts are compensated by a device (*34–38*). The present paper suggests that a clever use of local gravity fields could be useful to facilitate arm movements in different directions, depending on their acceleration or deceleration phase. Playing with local gravity fields could thus yield assistive control laws that are easily integrated by the human sensorimotor system. Therefore, locally varying gravity fields with a robotic exoskeleton may be an interesting approach for all applications where adapted weight support is relevant.

## Materials and Methods

In a series of 3 experiments, a total of 61 participants performed single-joint upward movements with their forearm in a parasagittal plane, with their upper arm resting in a vertical position as in previous protocols (*28, 48, 71*). The pointing movements thus consisted in point-to-point reaching movements with different gravitational torques induced at the elbow joint by a robotic exoskeleton.

### Experimental setup

#### Robotic exoskeleton

The experimental conditions described in the present study were achieved with the last actuated axis of the ABLE upper-limb exoskeleton as illustrated in Figure 2A. This exoskeleton has a total of four actuated joints. The first three joints correspond to the three main rotations of the human shoulder (abduction/adduction, internal/external rotation and flexion/extension) and were physically blocked in the present experiment. The last actuated joint corresponds to the flexion/extension of the human elbow. The forearm of the exoskeleton was equipped with a custom-made physical interface, previously developed and validated, minimizing unintended interaction efforts between the exoskeleton and the user (*73*).

The different gravity compensations were performed by means of a composite control law, based on an open-loop compensation of the exoskeleton dynamics (*72*) associated with both an individualized force feedback loop and feedforward terms resulting from a thorough population-based analysis (*48*).

#### Kinematic and electromyographic recording

An optoelectronic tracking system (10 Oqus 500+ cameras, 179 Hz; Qualisys, Gothenburg, Sweden) was used to record the position of nineteen 10 mm and one 3 mm reflective markers placed either on the participant or on the robot. Seven markers were placed on the participant: shoulder (acromion), elbow (epicondyle and epitrochlea), middle of the forearm, wrist (styloid process of the radius), base of the index proximal phalanx and tip of the index finger (the 3 mm marker). The other markers placed on the robot were used during the identification process.

The EMG activity was recorded with bipolar surface electrodes (Wave Plus, Wireless EMG, 2000 Hz; Cometa, Bareggio, Italy). The QTM interface (Qualisys, Gothenburg, Sweden) allowed recording synchronously kinematic data and EMG activity. Participants were first locally shaved and a hydroalcoholic solution was applied. The electrodes were then placed on the following four muscles: triceps (long and lateral heads), biceps brachii (long head) and brachioradialis. The EMG were placed according to the SENIAM recommendations (*82*).

#### Experimental task

For each participant, three targets indicated by LEDs were positioned in a parasagittal plane in front of the subject. The central target corresponded to a horizontal position of the forearm and the other two targets were distributed symmetrically with regard to a transverse plane as illustrated in Fig. 2A.

### Experiments 1 and 2

For all the experiments, written informed consent was given by each participant as required by the Helsinki declaration. The experimental protocols were approved by the local ethical committee for research (Université Paris-Saclay, 2017-34). All the involved participants were right-handed and did not have any known neurological or muscular disorders.

Twenty-two participants were involved in the Experiment 1(10 females; age: 24 ± 5 years; weight: 70.7 ± 9.7 kg; height: 176 ± 7.8 cm).

In Experiment 1, each participant was instructed to perform self-paced pointing movements toward semi-spherical targets of 2 – cm diameter. Participants were instructed that accuracy was not the primary concern of the task. The movement goal endpoint was the lit LED. The top and bottom LEDs were lit successively, which triggered 45° flexions and extensions of the human elbow ([−22.5°, 22.5°] centered on the horizontal). The LEDs were lit during 1.5 s and participants were instructed to complete each movement before they were switched off. This is significantly longer than the average duration previously observed for movements performed in same conditions with this exoskeleton (*72*), which ensured that participants could move at their preferred velocity. The experiment was divided in eighteen blocks of 15 trials. After a short familiarization with the task outside the exoskeleton, the identification of each participant’s anthropometric parameters was carried out following a preexisting protocol (*48*). This was necessary to accurately implement local, desired gravity fields. The participants were then asked to perform six blocks inside the exoskeleton in transparent mode, with minimized residual perturbations from the exoskeleton. This condition was referred to as 1g. The participants were then asked to perform six blocks inside the exoskeleton, either in mechanically induced zero-gravity (e.g. *g* ≈ 0 m.s^-2^) or in mechanically induced reversed gravity (e.g. g ≈ +9.81 m.s^-2^). These conditions were respectively referred to as 0g and −1g. The order of these two conditions was randomized. Between each block, two-minutes resting breaks were taken, during which the participants were asked not to move their forearm to avoid any readaptation to the Earth’s gravity field.

Twenty-nine participants were involved in the Experiment 2 (10 females; age: 23 ± 3 years; weight: 67.1 ± 11.8 kg; height: 175 ± 7.6 cm). Experiment 2 essentially followed the same protocol as Experiment 1. The differences were a different number of blocks in each gravity condition (e.g. 2 instead of 6) and a higher number of trials per block (e.g. 25 instead of 15).

### Experiment 3

Ten participants were involved in Experiment 3 (2 females; mean age, 24 ± 3 years; mean weight 69.9 ± 8.7 kg; mean height 176 ± 3.5 cm).

Experiment 3 globally followed the same procedure as Experiment 1. The specific characteristics of this experiment were: the blocks were composed of 15 elbow flexions as in Experiment 1 and each block was performed with a different mechanically induced gravity field chosen among 11 gravity fields with a ±0.2g increment. Half of the participants started the experiment in the 1g condition and were subjected to an increasing gravity compensation until they reached the −1g condition. The other half of the participants were subjected to the opposite gradient of gravity compensation: they started in the −1g condition and finished in the 1g condition. Between each block the same resting breaks as in Experiment 1 were taken. Experiment 3 was performed with a movement amplitude similar to Experiment 1 (e.g. 45°: [−22.5°, 22.5°] centered on the horizontal). Here, we decided to increment or decrement gravity continuously instead of randomly picking it among 11 possible values to favor the quickness of adaptation given that a restricted number of trials were recorded to limit the duration of the experiment and other effects like fatigue.

### Data processing and analysis

Kinematic and EMG data were processed using custom Python 3.8 scripts.

#### Kinematics

Three-dimensional position data of the marker taped on the tip of the index finger was used to assess the human kinematics. Data from the other reflective markers was used as a control. Position data was filtered (low-pass Butterworth, 5 Hz cutoff, fifth-order, zero-phase distortion, “butter” function from the “scipy” package) before differentiation as in previous studies (*28,48*). The computed kinematic parameters were defined as in Fig. 2B. The threshold for segmenting movements was set at 5% of PV in agreement with previous studies (*17,19,48*), which allowed defining MD and movement amplitude.

PV and PA were respectively defined as the maximum value of the velocity and the maximum positive value of the acceleration reached during each movement. In addition to these absolute parameters, the relative time to peak velocity (rtPV) was computed as the ratio between the time elapsed from the movement onset to PV and the duration of the movement.

A movement was considered invalid, and therefore removed, if the acceleration profile crossed 0 more than twice during the movement interval. This led to the exclusion of less than 1% of the movements performed by each participant.

#### EMGs

EMG data was first filtered (band-pass Butterworth, [20, 450] Hz cutoff, fourth order, zero-phase distortion, “butter” function from the “scipy” package), centered and rectified (*83*). Signals were normalized by the maximum peak value from all trials observed during the experiment for each participant and each muscle (*20, 48, 71*). The envelope of the signal was then obtained by filtering the pre-processed signal (low-pass Butterworth, 3 Hz cutoff, fifth order, “butter” function from the “scipy” package) (*83*).

A standard procedure was used to separate the tonic and phasic components of the pre-processed EMG signal (*20,34,84*). Averaged values of the integrated EMG signal were computed from 1 s to 0.5 s before the movement began and from 0.5 s to 1 s after the movement stopped, based on the velocity threshold. The segmentation process based on kinematics recording allowed to ensure that this baseline activity was not affected by the previous and upcoming movement. The tonic component of the EMG was computed as the linear interpolation between these two averaged values. The phasic component of the EMG was computed as the subtraction of the tonic component to the pre-processed signal.

Positive phasic components (referred to as activation), representing muscle bursts exceeding gravity-related EMG level, were assessed by computing the area of the positive phasic activity with a threshold set at 5% of the maximum measured value, which was reached during the acceleration phase. The negative phasic components (referred to as inactivation), were assessed by computing the absolute value of the area of negative phasic activity. A threshold set at −5% of the minimum (negative) measured value was used to detect these negative phasic components. As we dealt with upward movements (i.e. flexions), the activation of the flexors and the inactivation of the extensors were relevant to describe the impact of the various gravity fields on muscle patterns. These parameters were computed during the whole movement, without hypotheses on the beginning and end of activation/inactivation.

### Statistical analysis

Statistical analyses were performed on the averaged values of each participant in each block and/or condition. Normality (*Shapiro-Wilk* test (*85*)) and sphericity (*Mauchly’s* test (*86*)) of the distribution of the residuals were verified. Repeated measure analyses of variance (ANOVAs) were then performed between the different gravity conditions on the mean values obtained for each participant during Experiment 1 and Experiment 2. These analyses were used to account for differences in kinematics and muscle parameters. Significance of the ANOVA was corrected using a Greenhouse-Geisser method to correct sphericity issues (*ϵ* < 0.75). The significance level of the corrected *p*-value was set at *p* < 0.05.

Pairwise *t*-tests were used to perform post-hoc comparisons. These tests were corrected using Bonferroni method. The significance level of the corrected *p*-value was set at *p* < 0.05. All statistical analyses were performed using custom Python 3.8 scripts and the Pingouin package (*87*).

Given the number of participants and conditions in Experiment 3, only quantitative comparisons between the model predictions and the average experimental data were provided there. Mean absolute errors between the model and the experimental data for rtPV, and Pearson correlation coefficients were computed to quantify the agreement between the model and the data when varying gravity incrementally.

### Simulations

Movement kinematics were predicted using an optimal control model of the human forearm pointing task. Our model implemented the gravity-exploitation theory via the minimization of an effort-based cost considering the external action of the exoskeleton. To this aim, the Smooth-Effort model (SE) minimizing a compromise between the absolute work of the elbow net torque and the integrated squared jerk was tested (*9,28,88,89*). The dynamics of the human forearm when taking into account the exoskeleton were simulated as follows:

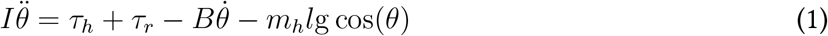

where *I* was the apparent inertia of the coupled human-exoskeleton system, *τ_h_* was the human net torque at the joint, *τ_r_* was the robot torque, *B* = 0.05 N.m.s.rad^-1^ was the damping of the human elbow (*90*) and the product *m*_h_^l^= 0.2934 kg.m, between the human mass and the weight moment arm, was identified following a preexisting procedure (*48*). The *I* = 0.35 kg.m^2^ term of the apparent inertia was an approximation of the total inertia of the human-exoskeleton system based on human dynamics identification, anthropometric tables and robot dynamics identification (*48, 72, 91, 92*). The robot torque was assumed to be known as it was accurately compensating for the human gravitational torque, which allowed rewriting Eq. 1 in the different gravity conditions as follows:

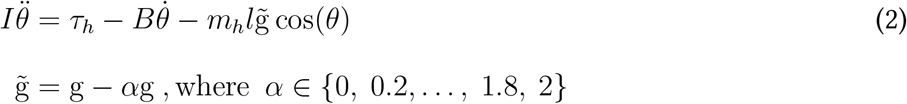

This equation could be differentiated to obtain the following expression for the angle jerk:

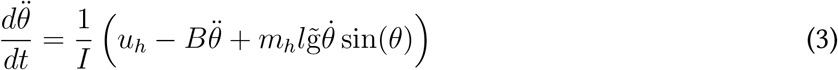

where 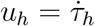 was the human control variable in a commanded torque change framework (*93*). In the chosen optimal control framework, motor planning is assumed to originate from the minimization of a cost function, expressed as follows:

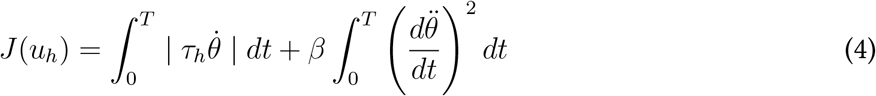

where *β* was a weight that allowed adjusting the importance given to each component.

All the simulation results reported in the present paper were obtained using the Matlab (MathWorks) version of “*gpops2*” (*94–96*). This software is based on an orthogonal collocation method and uses the “*SNOPT*” solver to solve the nonlinear programming problem (*97*).

## Acknowledgments

We thank Jérémie Gaveau for his comments on previous versions of the article. This work is supported by the “IDI 2017” project funded by the IDEX Paris-Saclay, ANR-11-IDEX-0003-02. This work is supported by the French National Agency for Research (grant ANR-19-CE33-0009).

## Author contributions

Conceptualization: DV, SB, FG, OB, NV, BB.

Methodology: DV, SB, NV, BB.

Investigation: DV, SB, BB.

Data processing: DV, SB.

Visualization: DV, BB.

Supervision: FG, OB, NV, BB.

Writing - original draft: DV.

Writing - review and editing: DV, SB, FG, OB, NV, BB.

## Data and materials availability

All data, code and materials used in the present study are available upon request to the corresponding author.

