## Supplementary for "Fast reoptimization of human motor patterns in non-Earth gravity fields locally induced by a robotic exoskeleton"

### Adaptation to 0g and -1g local gravity fields

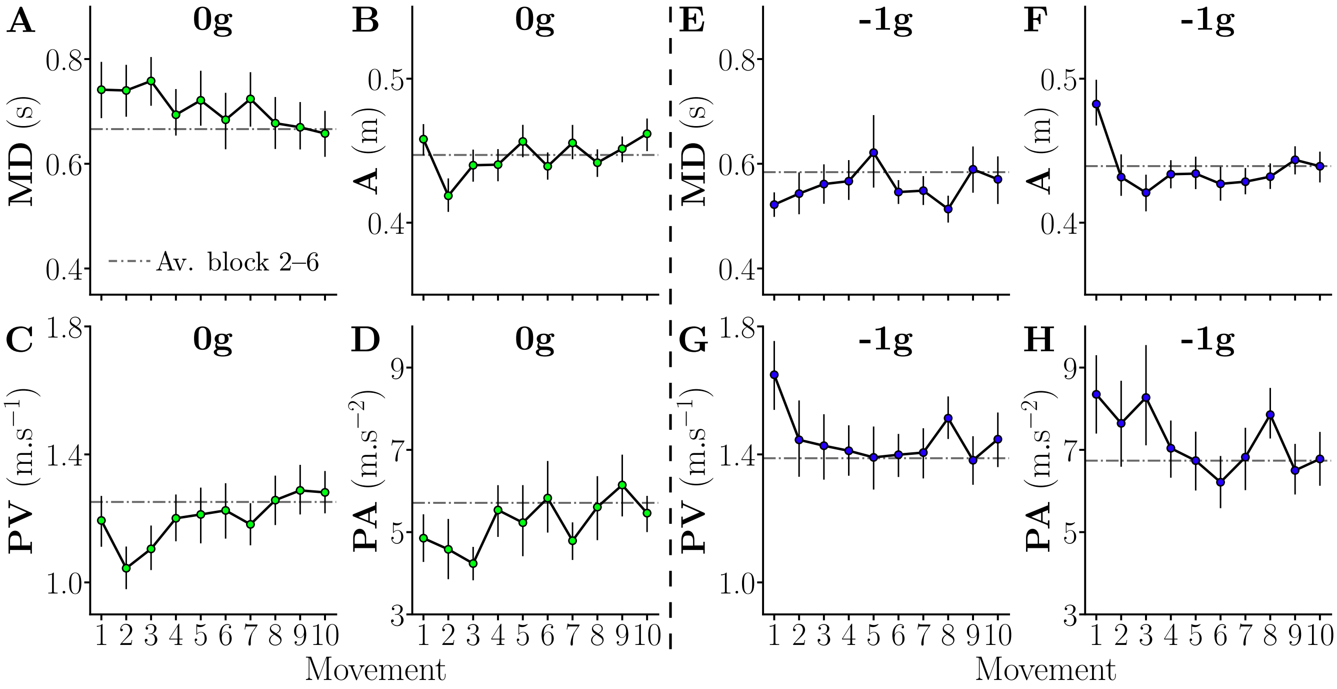


Figure S.1: Trends of adaptation during the first 10 movements in 0g and -1g, error bars represent the standard error across the 22 participants of Experiment 1. The adapted value of each parameter was computed as the average across blocks 2–6 and is emphasized by grey dotted lines. **A,E**. Movement duration MD. **B,F**. Amplitude A. **C,G**. Peak velocity PV. **D,H**. Peak Acceleration PA.

We analyzed the initial adaptation by pooling the 10 first movements from participants involved in Experiment 1 ($N = 22$). The adaptation to local 0g and -1g gravity fields induced by the exoskeleton was first analyzed on the basis of general kinematic parameters (see Fig. S.1). In the baseline 1g condition, the exoskeleton was set in transparent mode (*1, 2*), meaning that the participants were exposed to a normal gravity. Fast initial adaptation trends have already been reported on the same exoskeleton in the $1\text{g}$ condition (*3*).

We observed an initial adaptation in a couple of trials to the -$1\text{g}$ gravity field and no trends of adaptation to the $0\text{g}$ gravity field. The very first movements when exposed to reversed gravity were usually too fast and exhibited an overshoot (see Fig. S.1E–H). The overshoot amplitude was around $4\ \text{cm}$ for the first movement performed under reversed gravity. The lack of adaptation trends during the first movements performed under microgravity suggests a relatively accurate movement planning from information prior to movement onset.

These adaptation trends were statistically tested by performing pairwise $t$-tests between each movement and the adapted plateau obtained by computing the average value of each parameter across blocks $2$–$6$. In the $0\text{g}$ condition, these tests returned that the first movement was not significantly different from the adapted value in terms of $\text{MD}$ ($p = 0.19$), amplitude ($p = 0.34$), $\text{PV}$ ($p = 0.48$) and $\text{PA}$ ($p = 0.16$). In the -$1\text{g}$ condition, results were that only the first movement was significantly different from the adapted value in terms of $\text{MD}$ ($p = 0.024$), amplitude ($p = 0.029$) and $\text{PV}$ ($p = 0.047$), but not in terms of $\text{PA}$ ($p = 0.15$).

These results could be explained by the relatively strong upward force applied by the exoskeleton in the -$1\text{g}$ condition, which was probably underestimated by participants during motor planning. Such errors were quickly corrected in subsequent trials. In order to assess the origin of this overshoot from a motor command perspective, the initial adaptation was then analyzed through muscle activity based parameters (see Fig. S.2).

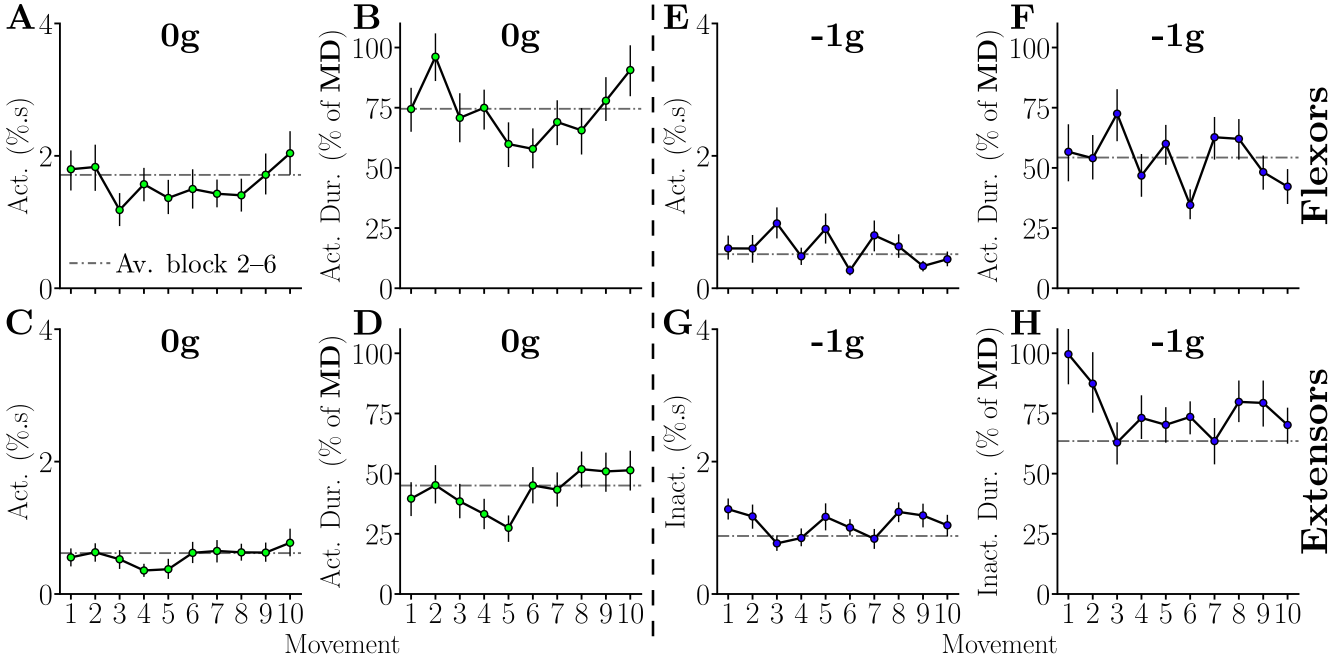


Figure S.2: Trends of adaptation during the first 10 movements in 0g and -1g, error bars represent the standard error across the 22 participants of Experiment 1. The adapted value of each parameter was computed as the average across blocks 2–6 and is emphasized by grey dotted lines. Flexors are illustrated on the upper line and extensors on the lower line. **A,E**. Flexors activation. **B,F**. Relative duration of flexors activation. **C**. Extensors activation. **D**. Relative duration of extensors activation. **G**. Extensors inactivation. **H**. Relative duration of extensor inactivation.

Interestingly, among all the analyzed parameters for both flexors and extensors (e.g. activation/inactivation, relative duration of activation/inactivation), only one parameters, computed in the -1g condition, exhibited a trend of adaptation (see Fig. S.2H). In the -1g condition, the too important relative duration of the inactivation of the extensors seemed to be responsible for the overshoot observed in the kinematic parameters.

These trends were statistically tested by following the same process as for kinematic parameters. Results in 0g were that the first movement was never significantly different from the plateau on any EMG based parameter ($p > 0.86$ and $p > 0.64$ for all flexors and extensors parameters respectively). Results in -1g were that the first movement was never different from the adapted plateau in terms of flexors behavior ($p > 0.62$ for all parameters). The extensors behavior during the first movement was significantly different from the adapted value ($p = 0.016$ and $p = 0.0079$ for inactivation and relative duration of inactivation respectively). From the second movement, no more significant differences were reported by the $t$-tests ($p > 0.067$ in all cases).

These results suggest that participants almost completely adapted their motor commands before the first movement onset, solely on the basis of haptic feedback while holding a static posture. The adaptation of the relative duration of extensors inactivation after the first movement in the -1g condition suggests that the static estimation led participants to underestimate the forthcoming interaction forces. This is consistent with gravity related efforts increasing during the acceleration phase of the tested movements. No clear trend of adaptation was observed for EMG parameters in the 0g condition, which is consistent with observations reported on kinematic parameters. Overall these results are consistent with previous works regarding the static estimation of forthcoming dynamics before movement onset (*4*).

Next, we investigated whether longer-term adaptation effects existed on the same kinematic parameters. Regarding EMG parameters, the adaptation was first assessed on the basis of flexors and extensors relative durations of activation and inactivation respectively. During Experiment 1, the participants performed a total of 90 upward movements (6 blocks $\times$ 15 trials) in each of the 1g, 0g and -1g conditions. The block-wise evolution of these parameters is depicted in Figure S.3. A visual inspection might suggest a potential adaptation for MD, PV and PA across blocks. However, when comparing the first and last blocks using Student's paired $t-$tests, we did not find any significant difference for all gravity conditions regarding all the parameters (MD: $p > 0.36$, A: $p > 0.68$, PV: $p > 0.44$, PA: $p > 0.51$, relative duration of flexors activation: $p > 0.24$ and relative duration of extensors inactivation: $p > 0.16$ for all gravity conditions).

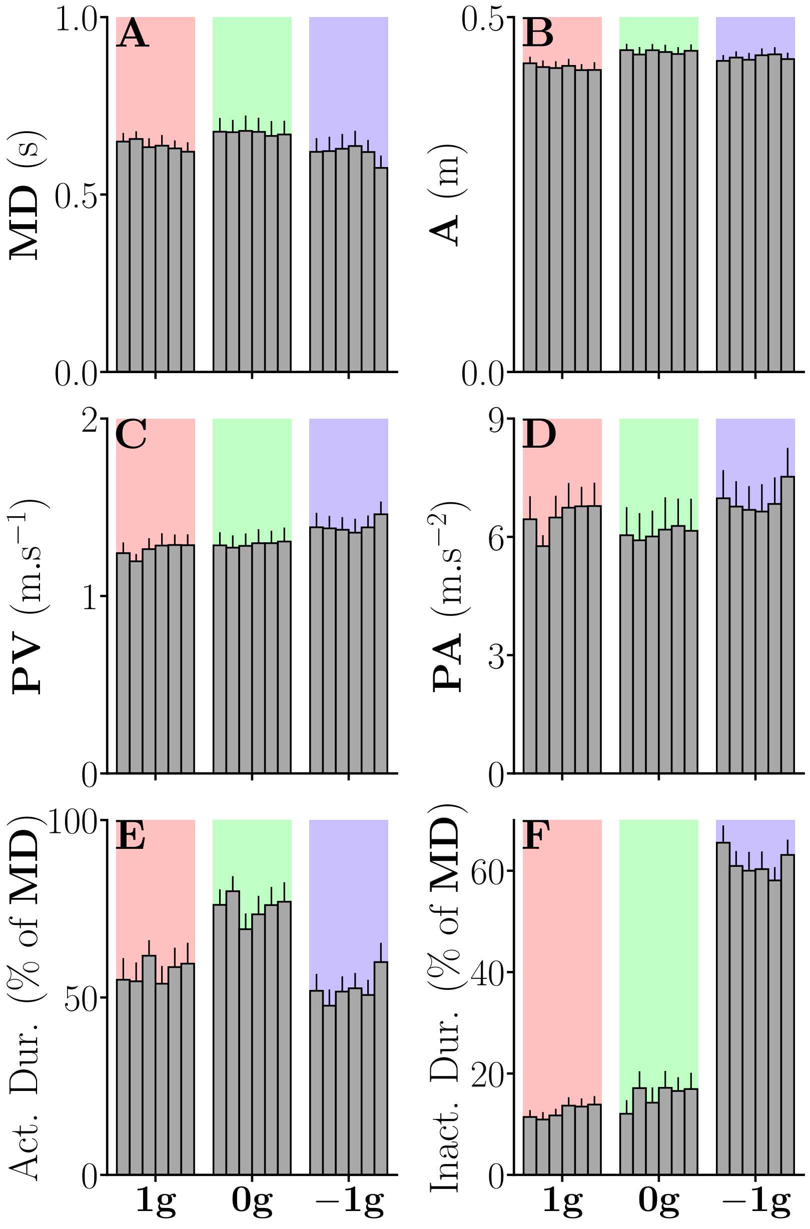


Figure S.3: Averaged parameters for each block, error bars represent the standard error across participants. **A**. Movement duration MD. **B**. Amplitude A. **C**. Peak velocity PV. **D**. Peak Acceleration PA. **E**. Relative duration of the flexors activation. **F**. Relative duration of the extensors inactivation.

The effect of gravity on movement kinematics being usually assessed from the rtPV and the activation and inactivation of flexors and extensors respectively (*5–11*), we also assessed the adaptation of these parameters (see Fig. S.4). Again, paired $t$-tests did not reveal any difference difference between the first and the last blocks of each gravity condition was found for the three parameters (rtPV: $p > 0.62$;

flexors activation: $p > 0.5$; extensors inactivation: $p > 0.23$ for all gravity conditions).

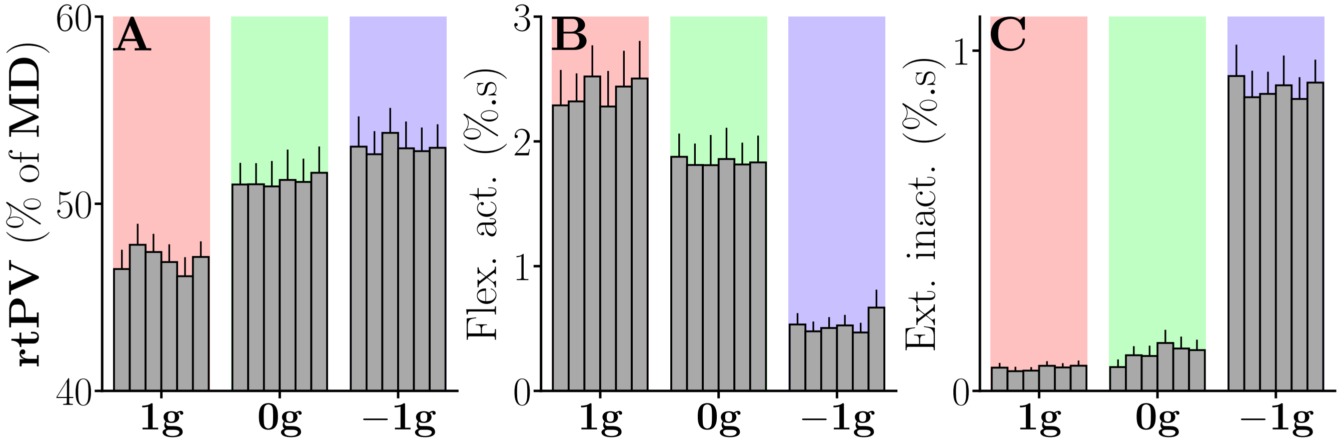


Figure S.4: Averaged parameters for each block, error bars represent the standard error across participants. **A**. Relative time to peak velocity rtPV. **B**. Flexors activation. **C**. Extensors inactivation.

All these analyses confirm that no major change in the forearm trajectories occurred across blocks and that a plateau was quickly attained. In the subsequent analyses, we thus averaged data across blocks and focused on the differences between gravity conditions.

3. Simon Bastide, Nicolas Vignais, Franck Geffard, and Bastien Berret. Analysing human-exoskeleton interaction: on the human adaptation to modified gravito-inertial dynamics. *Computer Methods in Biomechanics and Biomedical Engineering*, 22(sup1):S507–S509, may 2019.
